# The MPS1 kinase NTE region has helical propensity and preferred conformations towards the TPR domain

**DOI:** 10.1101/2020.03.31.018036

**Authors:** Yoshitaka Hiruma, Minos-Timotheos Matsoukas, Wouter G. Touw, Georgios Spyroulias, Geert J.P.L. Kops, Marcellus Ubbink, Anastassis Perrakis

## Abstract

The mitotic spindle assembly checkpoint (SAC) ensures accurate segregation of chromosomes by preventing onset of anaphase until all chromosomes are properly attached to spindle microtubules. The Monopolar spindle 1 (MPS1) kinase is one of the SAC components, localizing at unattached kinetochores by an N-terminal localization module. This module comprises a flexible NTE module and the TPR domain, which we previously characterized for their contribution to kinetochore binding. Here we discuss the conformations of the highly flexible NTE with respect to the TPR domain, using paramagnetic NMR. The distance restraints derived from paramagnetic relaxation enhancements (PREs) show that the mobile NTE can be found in proximity of a large but specific part of the surface area of the TPR domain. To sample the conformational space of the NTE in the context of the NTE-TPR module, we used the *ab initio* Rosetta approach supplemented by paramagnetic NMR restraints. We find that many NTE residues have a propensity to form helical structures and that the module localizes at the convex surface of the TPR domain. This work demonstrates the highly dynamic nature of the interactions between the NTE and TPR domains and it shows that the convex rather than the canonical concave TPR surface mediates interactions, leading to the auto-inhibition that the TPR exerts upon the NTE region in the context of SAC signaling.

## Introduction

Error free chromosome segregation is crucial to maintain genomic stability [1]. To ensure faithful segregation, a signaling cascade evolved to the spindle assembly checkpoint (SAC). The SAC monitors the attachment of spindle microtubules to kinetochores and prevents the onset of anaphase until all chromosomes are properly attached to spindle microtubules [2, 3]. The attachment of spindle microtubules is established with outer kinetochore components, in particular, the KNL1 complex and NDC80 complex [4]. When the NDC80 complex is not associated with spindle microtubules, MPS1 kinase initiates the SAC signaling by phosphorylating the KNL1 complex. MPS1, therefore, plays a major role to detect unattached kinetochore and has been recognized as a master regulator of SAC signaling [5]. MPS1 activity and turnover depend on the efficiency of kinetochore localization, which is mediated by three regions: an N-terminal extension (NTE); the adjacent tetratricopeptide repeat (TPR) domain; and the middle region (MR) further stabilizes the interaction of MPS1 to NDC80 complex [5]. The NTE motif is located at the N-terminus of MPS1 and is composed of sixty amino acids (Figure 1A). We have previously speculated that the interaction of the NTE module with the TPR exerts an auto-inhibitory function to the NTE, moderating the interaction with the NDC80C complex in the outer kinetochore [6]. Subsequently, an NMR analysis including the assignment of most amide resonances (87%) of the NTE-TPR module, showed that the NTE does not adopt a defined structure, except for a short segment (residues, 14-23), which has a propensity to form a helical turn [7]. This helical fragment is required for interactions with kinetochores and forms intermolecular interactions with the TPR domain. Notably, bypassing this NTE-TPR interaction results in high MPS1 levels at kinetochores, inefficient MPS1 delocalization upon microtubule attachment, and SAC silencing defects [7]. As it became clear that perturbations of the NTE lead to notable physiological changes in the dynamics of the MPS1-NDC80C interactions and mitotic defects, we wanted to further study the dynamics of the NTE. However, solving a solution structure of the NTE with conventional methods was considered impossible, as all data suggested it is very flexible, essentially behaving as an intrinsically disordered region (IDR).

**Figure 1.**
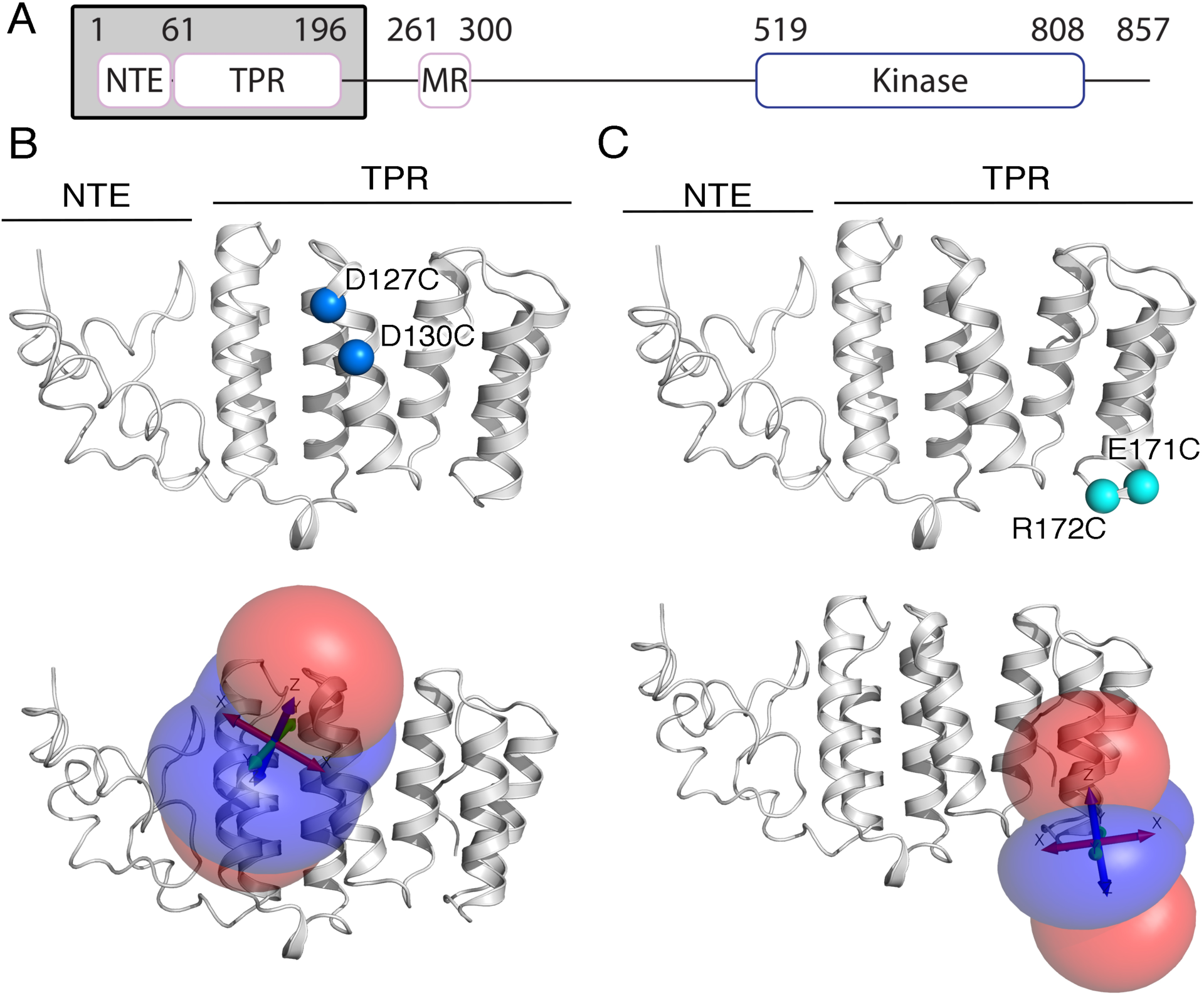
A) Schematic diagram of Mps1. B) Double-cysteine mutation position (D127C/D130C shown in blue) for lanthanoid tag attachment via the cysteine side chains that were used to measure PCSs and isosurfaces representing the Δχ tensor for the double-armed Yb^3+^ corresponding to a PCS of ±0.5 ppm. C) Double-cysteine mutation position (E171C/R172C shown in cyan) for lanthanoid tag attachment and isosurfaces representing the Δχ tensor for the double-armed Yb^3+^ corresponding to a PCS of ±0.5 ppm. Positive and negative PCS values are indicated by blue and red, respectively. The figure was prepared using Numbat [22].

Structural understanding of the conformations of such regions can be gained by paramagnetic NMR techniques. In particular, paramagnetic relaxation enhancements (PREs) have proven to be a useful tool to characterize the dynamic nature of proteins [8]. By introducing paramagnetic centers at specific sites of a protein, PREs can be observed for the surrounding nuclear spins. The PRE effect has a strong distance dependence (*r*^-6^). As the averaged relaxation rate will be dominated by nuclei in close proximity of the paramagnetic center, sparsely populated states of protein conformations can be detected. Pseudocontact shifts (PCSs) provide independent information about the averaged position of a nucleus relative to the paramagnetic center. Its distance dependence is less strong (r^-3^) and the PCS is also angle dependent. The PCS is much less influenced by minor states. Thus, PRE and PCS provide complementary information about protein ensembles [8]. It should be noted that that the structural characterization of ensembles of conformations is an ill-posed inverse problem since the NMR observables are ensemble averages. Thus, there is an infinite number of ensembles of structures that can satisfy the experimental data equally well [9]. Additional modeling is required to better characterize the conformational space. Various approaches have been developed over the past decades [10–13]. Maximum Occurrence analysis, for instance, provides the maximum percent of time that a conformer can exist by assigning a value for each conformer [9]. Other approaches involve the use of the *ab initio* Rosetta protocol method coupled with paramagnetic data [14–16].

Here, we report a structural characterization of the NTE module and its interaction with the TPR module, based on distance restraints obtained from PREs and PCSs. Two sets of lanthanoid tags and spin labels were attached at specific sites of the NTE and TPR domains and PREs and PCSs were measured. The solution structures represented by an ensemble of 12 structural models indicate large conformational freedom of the NTE module, as expected for an IDR. Upon interacting with the TPR domain, the NTE has a helical propensity and the interactions occur selectively on one side of the TPR domain, helping to understand the dynamic turnover of MPS1 kinetochore localization.

## Materials and methods

### Chemicals

^15^NH_4_Cl, and D_2_O were purchased from CortecNet. CLaNP-7 (caged lanthanoid NMR probe, version 7) was synthesized and loaded with diamagnetic lutetium ion (Lu^3+^) or paramagnetic ytterbium ion (Yb^3+^) as previously described [17]. MTS (1-acetyl-2,2,5,5-tetramethyl-3-pyrroline-3-methyl)-methanethiosulfonate) and MTSL were purchased from Toronto Research Chemicals.

### Paramagnetic NMR probe attachment

All proteins were expressed and purified as described in [7]. First, endogenous Cys residues, Cys139 and Cys229, were substituted to Ser and Ala, respectively, to prevent unspecific attachment of the probes. In addition, Asp127 was substituted to Ala to enhance the yield of the MPS1 variants. Site-directed mutagenesis was performed using the QuikChange protocol (Stratagene). To measure PCSs, we used CLaNP-7, which has been demonstrated to be a useful tool for the structure determination [17, 18]. CLaNP-7 covalently attaches to two Cys residues in a bidentate manner, which ensures a minimal mobility of the probe on the protein surface. Based on the previous analysis of chemical shift perturbations [7], two pairs of Cys residues were introduced on the TPR domain, D127C/D130C (construct A) and E171C/R172C (construct B). These two double Cys mutants of MPS1 constructs (residues 1-196) were used for CLaNP-7 attachment. Concurrently, two single Cys mutants of MPS1 constructs (residues 1-239), I23C (construct C) and S42C (construct D), were prepared for MTS/MTSL tagging. The MPS1 variants were linked to the paramagnetic NMR probe as follows. The purified protein samples were incubated with freshly prepared 3 mM DTT in buffer A (20 mM KPi, pH 7.5), supplemented with 150 mM KCl for 30 minutes on ice. The samples were then loaded on Superdex G75 16/60 HiLoad (GE Healthcare) pre-equilibrated in buffer A supplemented with 150 mM KCl. The protein fractions were pooled and mixed with three molar equivalents of the paramagnetic NMR probes and incubated at 4 °C overnight. The samples were two-fold diluted in buffer A with 50 mM KCl and loaded on a HiTrap Heparin HP column (GE Healthcare). After washing with the same buffer, the protein was eluted in buffer A containing 500 mM KCl. The sample was then loaded on a Superdex G75 16/60 HiLoad (GE Healthcare) pre-equilibrated in the NMR buffer (20 mM HEPES/NaOH, pH 7.4, 150 mM KCl and 10% D_2_O). The protein fractions were pooled and concentrated.

### NMR measurements, assignment and data analysis

NMR samples contained 100-500 µM MPS1 variants labeled with Ln^3+^-CLaNP-7/MTS/MTSL. Two-dimensional ^15^N-^1^H TROSY/HSQC spectra [19] were recorded at 298 K on a Bruker AVIIIHD 850 spectrometer equipped with a TCI-Z-GRAD cryoprobe.

All NMR data were processed in Topspin 3.1 (Bruker, Biospin) and analyzed in CCPNMR [20]. The assignments for amide resonances of MPS1 (residues 1-239 and 62-239) were based on previous work (BMRB entry, 27641 and 27642) [7].

The location of the Ln^3+^ ion and Δχ-tensor were calculated with Numbat [22] using the crystal structure of the MPS1 TPR domain (PDB entry, 4H7Y) [21] and the PCSs for residues 59-199. The quality (Qa) scores of PCSs were calculated as previously described [23], using the following formula.

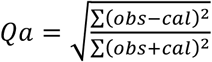

PRE datasets were evaluated in a manner similar to what was previously reported [24]. The peak intensity of amide resonances in the spectra of the paramagnetic (MTSL) and diamagnetic (MTS) samples were represented as *I*_para_ and *I*_dia_, respectively [25]. The intensity ratios of *I*_para_ / *I*_dia_ were normalized as described previously. The R_2,para_, was calculated and converted to distances as previously reported [25].

### Rosetta calculations

Using sequence information from the target protein, the Rosetta *ab initio* sampling algorithm was used, utilizing the Monte-Carlo assembly of nine and three residue fragments [14]. The fragment libraries for MPS1 (residues 1-199) were obtained from the Robetta server [26] with the additional input of backbone atom diamagnetic chemical shift data for the structure, obtained from the NMR measurements for high-quality generated fragments. For the construction of a full set of coordinates for the starting model of MPS1 NTE and TPR domains, the X-ray crystal structure (PDB ID: 4B94) [6], which does not provide coordinates for the NTE domain, was used as a template to add missing residues for the entire NTE-TPR domain (residues 1-199) with the program Modeller, using default settings [27].

Initially, 1,000 decoy structures were generated, setting the coordinates of the region containing residues 60-199 as rigid, and without any PCS restraints. The generated decoys were scored against PCS data with unity weights. Then, the PCS fit-quality scores for each of the two metal centers were independently weighted relative to the Rosetta low-resolution scoring function and used in the subsequent modeling round as independent scoring factors for each metal center, as described previously [15].

Next, 10,000 all-atom models were generated in Rosetta calculations. Again, the coordinates of the region containing residues 60-199 were considered rigid, but PCS data were used from the two metal binding tags, directing the Rosetta’s conformational sampling to satisfy the input PCS restraints, as PCS restraints from multiple metal binding tags are expected to dramatically improve the sampling process [28]. We utilized three classes of intermolecular PRE distance restraints obtained from the ^15^N-enriched MPS1 constructs I23C and S42C, linked to MTSL (Table 1). Residues strongly affected by MTSL and whose resonances disappeared in the paramagnetic spectrum were restrained with only an upper bound of 13 Å, residues affected by MTSL were restrained only with a lower limit of 22 Å and residues affected by the spin label and whose resonances are observed in the paramagnetic spectra were restrained with both upper and lower bounds as previously described [29]. All restraints were implemented as distances between the CB atom of the tagged residue and the N atom of the corresponding residue. Margins of 4 Å were used for all the calculated restraints to accommodate the experimental error and enhance sampling [30]. The all-atom models generated, were subsequently scored with Rosetta’s all-atom energy function [31] and weighted PCS energy for each of the two metal tags.

**Table 1.**
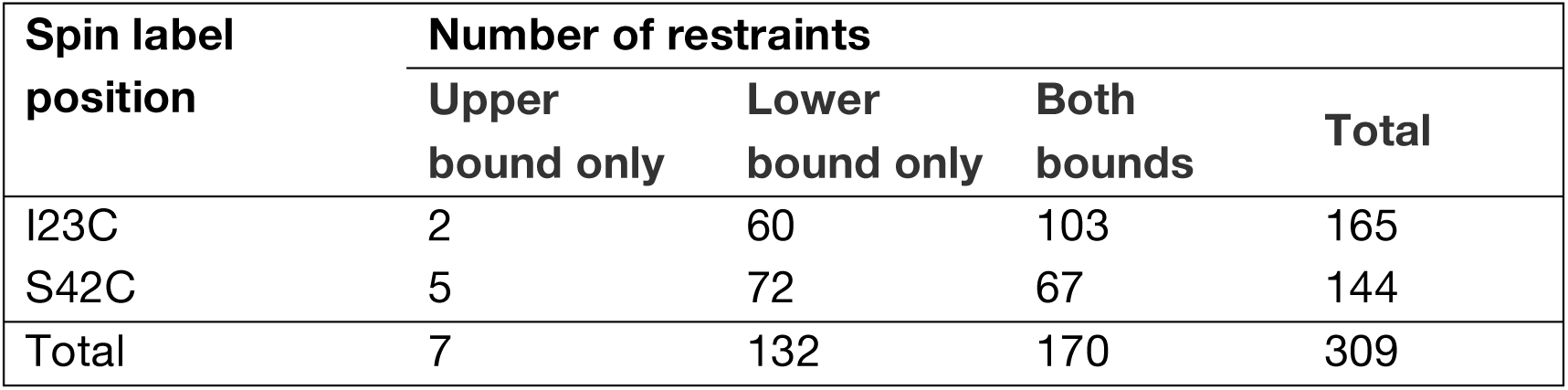
Intermolecular PRE distance restraints used in Rosetta calculations.

| Spin label position | Number of restraints |  |  | Total |
| --- | --- | --- | --- | --- |
|  | Upper bound only | Lower bound only | Both bounds |  |
| I23C | 2 | 60 | 103 | 165 |
| S42C | 5 | 72 | 67 | 144 |
| Total | 7 | 132 | 170 | 309 |

### Computational analysis

Secondary structure analysis of the ensemble of generated conformations was performed using DSSP [33] from the GROMACS 2018.5 tools [32] which was also used for other types analysis. Clustering of the final 1,000 models was performed using the FCC clustering algorithm, with which a matrix based on common contacts between NTE and TPR was calculated [34]. The similarity matrix was set to 0.5 and the minimum cluster size was set to 6, based on software default values and manual tuning.

## Results and Discussion

To determine the dynamic structure of the NTE module with respect to the TPR domain, a set of distance and orientation restraints were obtained from paramagnetic NMR experiments. To investigate the relative position of the NTE to the TPR module, the paramagnetic CLaNP-7 probe was covalently attached to two Cys residues in a bidentate manner. As shown in Figure 1B, MPS1 D127C/D130C (construct A) and E171C/R172C (construct B) were designed. The ^15^N labeled constructs were linked to Lu^3+^ or Yb^3+^ loaded CLaNP-7 and two-dimensional heteronuclear correlation spectra were recorded (Supplementary Figure 1A). Due to lack of information for the NTE structure, only PCSs measured for the TPR domain were used for the initial analysis. PCSs derived distance and orientation restraints were used to calculate the position of the lanthanoid ion (Ln^3+^) and orientation of principle components of the anisotropy of the magnetic susceptibility anisotropy (ΔX) tensor (Figures 1C,D, Supplementary Table 1). These parameters are required to convert PCSs to accurate restraints for structure determination. Having established the attachment site of CLaNP-7, we measured PCSs of the NTE module. The derived distance and orientation restraints were used for the subsequent structure calculation. In parallel, to investigate the surface area of the TPR domain being sampled by the NTE, an MTSL spin label was attached to two positions of the NTE. ^15^N-enriched MPS1 I23C (construct C) and S42C (construct D) were linked to MTSL or MTS and PREs were measured (Supplementary Figure 1B). In comparison to PCS distances, which are generally in good agreement with those calculated from the crystal structure, PRE derived distances can reflect transient states in populations as low as 0.5% [35]. In total, 501 restraints were obtained; two sets of PREs derived distances from the NTE (309 restraints) and two sets of PCSs derived distances (192) and orientations from the TPR.

To characterize the ensemble conformation of the NTE module and its interaction with the TPR domain, an ensemble approach was employed using the paramagnetic data as restraints. Structure calculations were carried out with the Rosetta *ab initio* sampling algorithm utilizing the Monte-Carlo assembly of nine and three residue fragments, obtained from the Rosetta server, using backbone atom diamagnetic chemical shift data from the NMR measurements. The 10,000 generated all-atom models, which were calculated using the PCS and PRE restraints, were rescored with Rosetta’s energy function and the weighted PCS energy for the two metal tags (see Methods). The 1,000 structures with the lowest combined PCS and Rosetta energy terms, were selected for further analysis. To understand the agreement of these models with the experimental PRE restraints, we calculated for each model the cumulative PRE restrain violation (the sum of all violations for each model, Figure 2A, B). Further we examined the portion of the models that show extreme violations for each individual PRE restraint; 95% of the individual PRE distance restraints are satisfied by the average back-calculated distance from the models (Figure 2C). Further analysis on the secondary structure of the final 1,000 models, demonstrated that NTE regions with residues 3-5, 11-16, 49-56 showed more than 50% propensity of forming α-helical segments (Figure 2D). To gain a better understanding of contacts formed between NTE and the TPR domain, average distances were calculated from the ensemble of the final 1,000 models (Figure 2E). Region 15-25 is frequently in proximity to residues 85-100 (helix α2) and less frequently to residues 110-135 (helices α3, α4 and their connecting loop), while residues 40-45 of the NTE interact with 85-95 (helix α2). Strikingly, helix α1 of the TPR domain, to which the NTE is connected, is poorly represented in the contact matrix, a fact that highlights the tendency of NTE to interact mostly with α2, α3 and α4. To further understand the distribution of conformations within the ensemble of the 1,000 lowest energy structures, we performed a clustering on the basis of the fraction of common contacts between the NTE and TPR domains, to obtain twelve total clusters; the lowest combined-energy model from each cluster is shown in Figure 3A, as a cluster representative.

**Figure 2.**
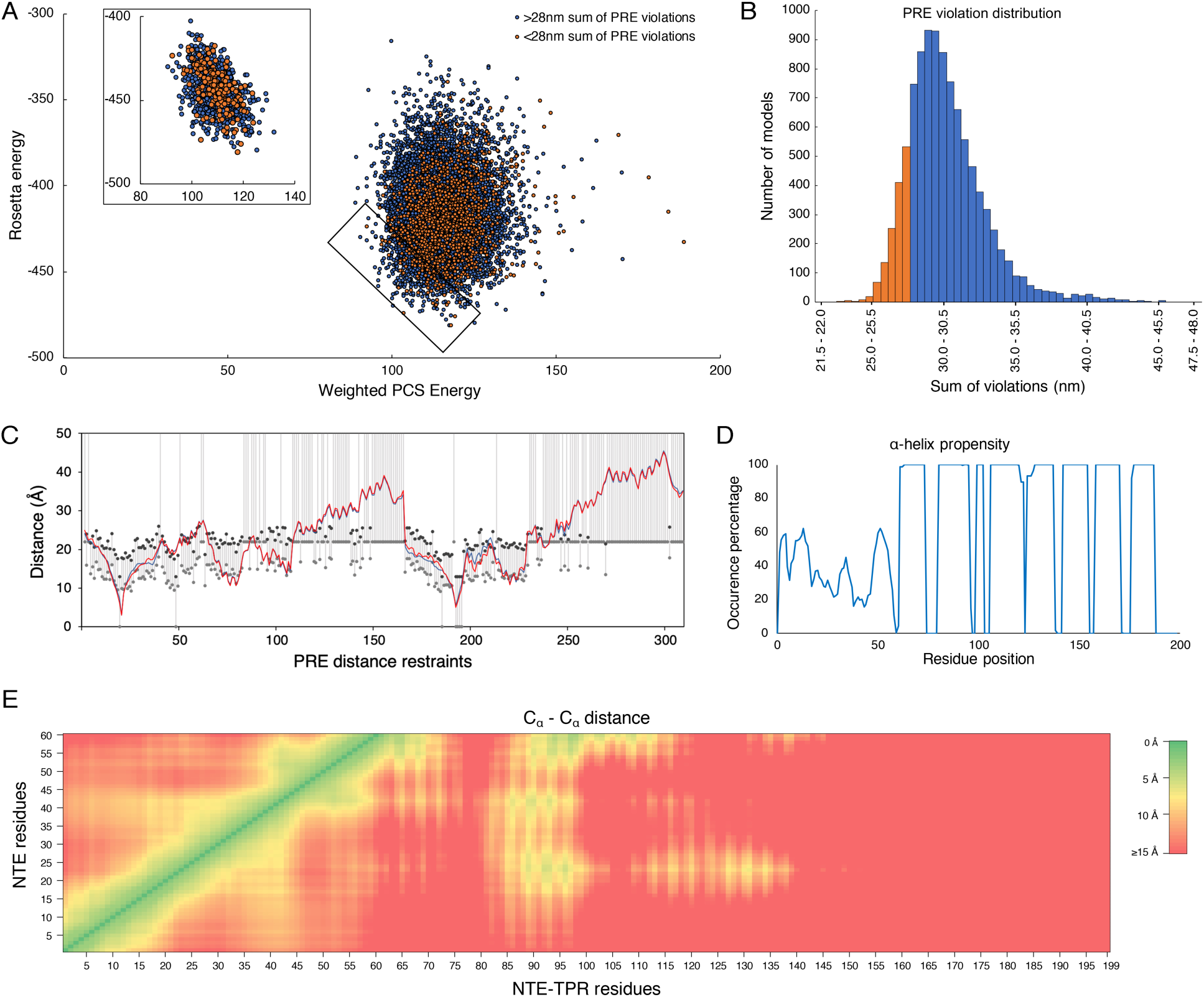
A) Rosetta vs weighted PCS energy of the ensemble of 10,000 structures. Orange points correspond to models where the cumulative PRE distance restraint violation is less than 28 nm, chosen to visualize the 15% of models that agree better with the experimental restraints; blue points represent the other 85% of models with a larger cumulative restraint violation. The inset depicts the 1,000 lowest combined energy models used for further analysis. B) Distribution of models over and under the 28 nm PRE violation threshold. C) Back-calculation of distances corresponding to each PRE derived distance restraint. Lower and upper limits of distance restraints are shown in grey and black dots respectively, interconnected by grey lines. Blue and red lines depict average distances for each distance restraint for the final 1,000 and the representative 12 models, respectively. D) α-helix propensity for each residue on the 1000 low energy models. E) Distance matrix illustrating Cα – Cα average distances from the 1000 lowest combined energy models between NTE and the rest of the domain residues.

**Figure 3.**
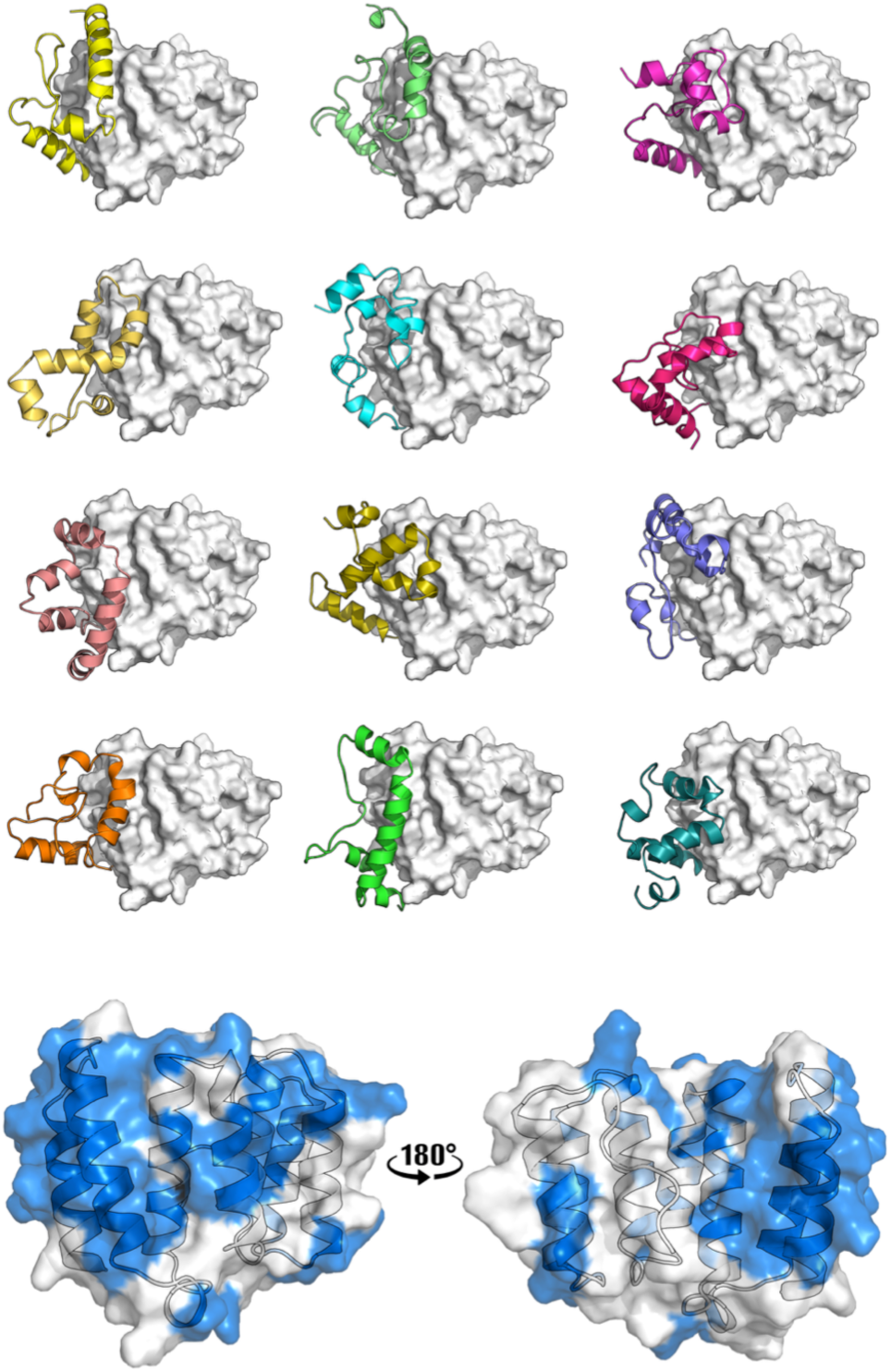
A) A representative model with the lowest energy selected from each of the 12 clusters B) The surface of the TPR coloured by the observed PREs.

The current ensemble of structures confirms that despite its flexibility, the NTE interaction is dominated by interactions with the proximal side of the convex outer surface of the TPR domain, showing also the strongest PRE effects (Figure 3B). In principle, the size of the NTE is sufficient to span across the inner surface of the convex of the TPR domain. However, two of the N-terminal α-helices of the TPR domain consist of multiple basic residues (Lys 71, Lys 74, Lys 86 and Arg 90), which likely restrict the conformational space of the negatively charged NTE, in line with our previous work showing that the NTE-TPR interaction depends largely on electrostatic interactions. Previously, we have suggested, based on analysis of NMR spectra [7], that the region between residues 14-23 has a high propensity to form a helix. Disrupting that helix formation by substituting Asn18 with Pro resulted in loss of the kinetochore localization of MPS1 [7]. The Rosetta modeling approach here, shows this region to participate in a helical structure in 24-52% of the models, while the regions 3-5, 11-16, 49-56 participate in helices in more than 50% of the models. This propensity might well hint toward an induced fit event, where the helix forms upon interaction with the NDC80C in the outer kinetochore.

The MPS1-NDC80C interaction has been reported to be facilitated by NTE phosphorylation [36, 37]. MPS1 is heavily phosphorylated in mitotic cells and there are eight reported phosphorylation sites in the NTE [21]. Despite some attempts we could not obtain a homogenous population of phosphorylated NTE-TPR module to compare the dynamics in the presence of phosphorylation (Supplementary Figure 1C). Preliminary comparison analysis show that the largest chemical shift perturbations were observed at the NTE residues 40-60. Recently, phosphorylation of the NTE residues 40-49 in the NTE has been reported to play essential roles in dimerization and activation of the MPS1 kinase. MPS1 requires auto-transphosphorylation on its kinase domain to be fully catalytically competent and the NTE has been believed to be involved in the regulation of the dimerization and transphosphorylation [38]. Previously, the MPS1 NTE-TPR module was indicated to form homodimer [21]. With the current structures, we can conclude that the NTE-TPR domain does not mediate MPS1 dimerization *per se*. However, we cannot exclude a possibility that MPS1 forms dimer through the interaction of the NTE with other domains, as recently proposed by Combes *et al. [39]*.

Overall, we have used paramagnetic data together with the *ab initio* Rosetta algorithm, and further computational analysis, to explore the NTE movement and dynamics in respect to the TPR domain. The initial chemical shifts, assisted the fragment selection process for *ab initio* modelling in Rosetta. The hybrid methodology we implemented here, also involved the usage of paramagnetic NMR data (PCS and PRE) restrains. This approach can be of general use in modelling the interactions of IDRs and folded domains of known structures. Further, we propose that the contact-based clustering is useful for selecting representative structural models in the context of this workflow.

In conclusion, we demonstrate an ensemble of NTE conformations that associate and dissociate dynamically with the TPR surface and are likely associated with the previously described auto-inhibitory action exerted by the TPR towards NTE-mediated interactions. The data indicate a propensity for the NTE in forming helical segments that might hint toward an induced fit mechanism mediating NTE interaction with the outer kinetochore components.

## Acknowledgements

We would like to thank Dr. Kala Bharath Pilla for constructive discussions and guidance on analyzing NMR data and performing Rosetta calculations.

## Supplementary materials

**Supplementary Figure 1.**
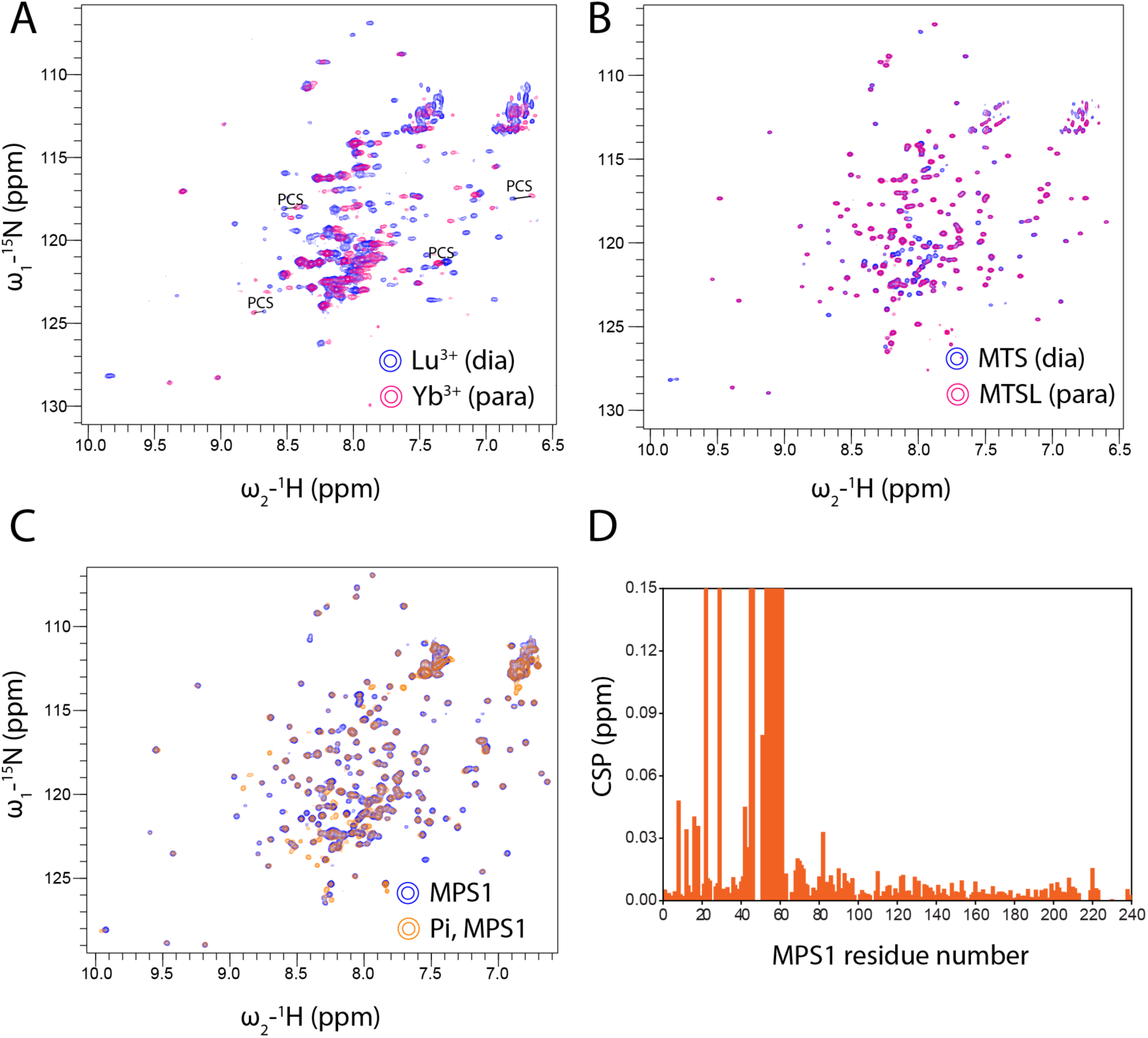
A) TROSY/HSQC spectra of Ln^3+^-CLaNP-7 labeled MPS1. Blue and magenta peaks represent Lu^3+^ and Yb^3+^ containing samples, respectively. The PCS are illustrated for several residues by solid lines. B) MTS/MTSL tagged MPS1. C) Phosphorylated samples of MPS1. D) CSP analysis of MPS1 samples with and without phosphorylation.

**Supplementary Figure 2.**
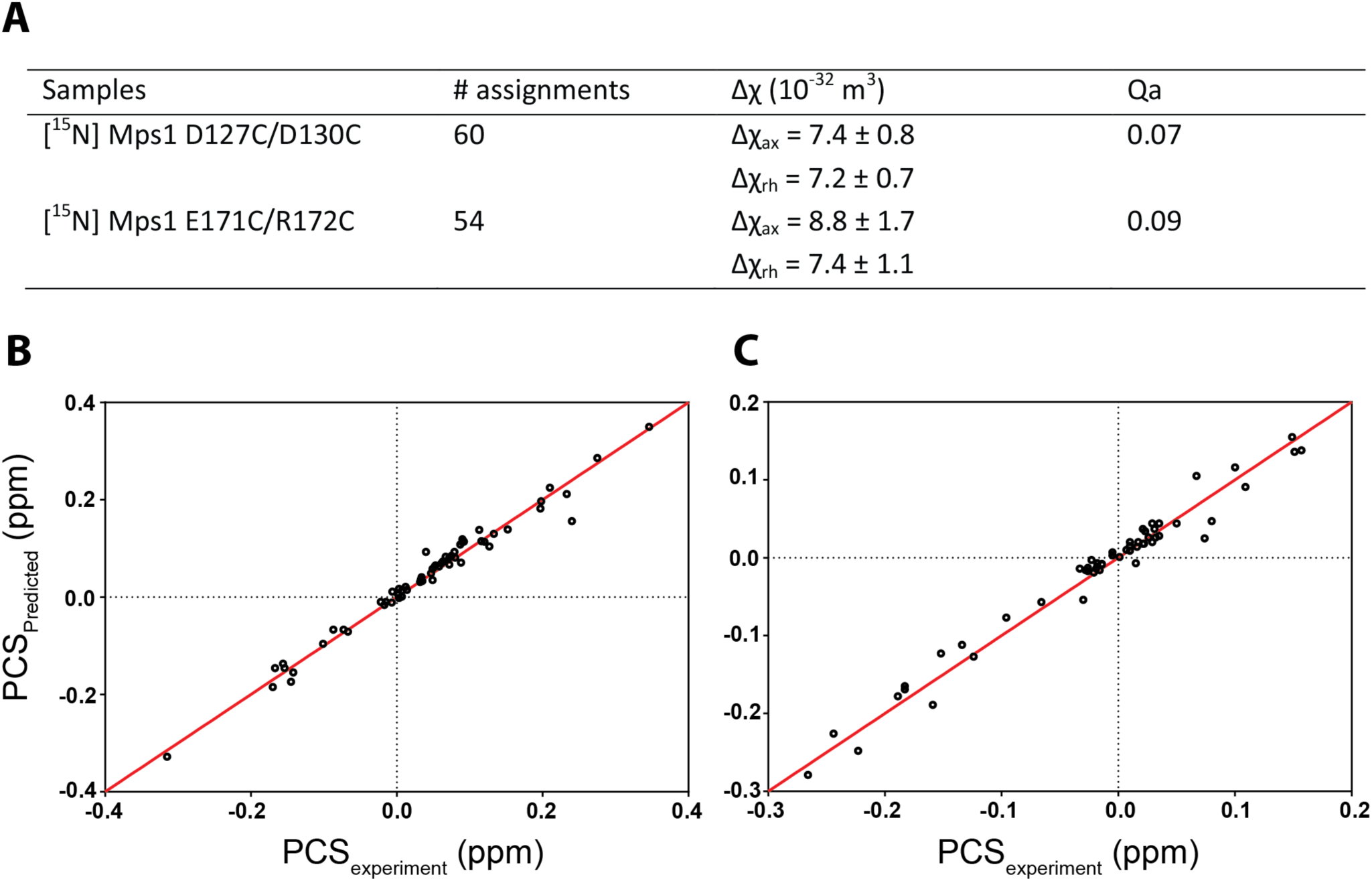
Yb^3+^-CLaNP7 was used as a paramagnetic center. A) Summary of PCS analysis. B-C) Plots of back-predicted PCSs versus experimentally measured PCSs. The ideal correlation is indicated by the solid red line. B) D127C/D130C construct. C) E171C/R172C construct.

## References

1. Ricke RM, Van Deursen JM (2013) Aneuploidy in health, disease, and aging. J. Cell Biol.

2. Etemad B, Kops GJPL (2016) Attachment issues: Kinetochore transformations and spindle checkpoint silencing. Curr. Opin. Cell Biol.

3. London N, Biggins S (2014) Signalling dynamics in the spindle checkpoint response. Nat. Rev. Mol. Cell Biol.

4. Musacchio A, Desai A (2017) A Molecular View of Kinetochore Assembly and Function. Biology (Basel). https://doi.org/10.3390/biology6010005

5. Pachis ST, Kops GJPL (2018) Leader of the SAC: Molecular mechanisms of Mps1/TTK regulation in mitosis. Open Biol.

6. Nijenhuis W, Von Castelmur E, Littler D, et al (2013) A TPR domain-containing N-terminal module of MPS1 is required for its kinetochore localization by Aurora B. J Cell Biol. https://doi.org/10.1083/jcb.201210033

7. Pachis ST, Hiruma Y, Tromer EC, et al (2019) Interactions between N-terminal Modules in MPS1 Enable Spindle Checkpoint Silencing. Cell Rep. https://doi.org/10.1016/j.celrep.2019.01.017

8. Hass MAS, Ubbink M (2014) Structure determination of protein-protein complexes with longrange anisotropic paramagnetic NMR restraints. Curr. Opin. Struct. Biol.

9. Carlon A, Ravera E, Andralojc W, et al (2016) How to tackle protein structural data from solution and solid state: An integrated approach. Prog. Nucl. Magn. Reson. Spectrosc.

10. Chen JL, Wang X, Yang F, et al (2016) 3D Structure Determination of an Unstable Transient Enzyme Intermediate by Paramagnetic NMR Spectroscopy. Angew Chemie - Int Ed. https://doi.org/10.1002/anie.201606223

11. Camilloni C, Vendruscolo M (2015) Using Pseudocontact Shifts and Residual Dipolar Couplings as Exact NMR Restraints for the Determination of Protein Structural Ensembles. Biochemistry. https://doi.org/10.1021/acs.biochem.5b01138

12. Salmon L, Bascom G, Andricioaei I, Al-Hashimi HM (2013) A general method for constructing atomic-resolution RNA ensembles using NMR residual dipolar couplings: The basis for interhelical motions revealed. J Am Chem Soc. https://doi.org/10.1021/ja400920w

13. Berlin K, Castañeda CA, Schneidman-Duhovny D, et al (2013) Recovering a representative conformational ensemble from underdetermined macromolecular structural data. J Am Chem Soc. https://doi.org/10.1021/ja4083717

14. Schmitz C, Vernon R, Otting G, et al (2012) Protein structure determination from pseudocontact shifts using ROSETTA. J Mol Biol. https://doi.org/10.1016/j.jmb.2011.12.056

15. Pilla KB, Leman JK, Otting G, Huber T (2015) Capturing conformational states in proteins using sparse paramagnetic NMR data. PLoS One. https://doi.org/10.1371/journal.pone.0127053

16. Gu L, Tran J, Jiang L, Guo Z (2016) A new structural model of Alzheimer’s Aβ42 fibrils based on electron paramagnetic resonance data and Rosetta modeling. J Struct Biol. https://doi.org/10.1016/j.jsb.2016.01.013

17. Liu WM, Keizers PHJ, Hass MAS, et al (2012) A pH-sensitive, colorful, lanthanide-chelating paramagnetic NMR probe. J Am Chem Soc. https://doi.org/10.1021/ja307824e

18. Hiruma Y, Hass MAS, Kikui Y, et al (2013) The structure of the cytochrome P450cam-putidaredoxin complex determined by paramagnetic NMR spectroscopy and crystallography. J Mol Biol. https://doi.org/10.1016/j.jmb.2013.07.006

19. Pervushin K, Riek R, Wider G, Wuthrich K (1997) Attenuated T2 relaxation by mutual cancellation of dipole-dipole coupling and chemical shift anisotropy indicates an avenue to NMR structures of very large biological macromolecules in solution. Proc Natl Acad Sci. https://doi.org/10.1073/pnas.94.23.12366

20. Vranken WF, Boucher W, Stevens TJ, et al (2005) The CCPN data model for NMR spectroscopy: Development of a software pipeline. Proteins Struct Funct Genet 59:687–696. https://doi.org/10.1002/prot.20449

21. Thebault P, Chirgadze DY, Dou Z, et al (2012) Structural and functional insights into the role of the N-terminal Mps1 TPR domain in the SAC (spindle assembly checkpoint). Biochem J. https://doi.org/10.1042/BJ20121448

22. Schmitz C, Stanton-Cook MJ, Su XC, et al (2008) Numbat: An interactive software tool for fitting δχ-tensors to molecular coordinates using pseudocontact shifts. J Biomol NMR. https://doi.org/10.1007/s10858-008-9249-z

23. Lescanne M, Ahuja P, Blok A, et al (2018) Methyl group reorientation under ligand binding probed by pseudocontact shifts. J Biomol NMR. https://doi.org/10.1007/s10858-018-0190-5

24. Schilder J, Löhr F, Schwalbe H, Ubbink M (2014) The cytochrome *c* peroxidase and cytochrome *c* encounter complex: The other side of the story. FEBS Lett 588:1873–1878. https://doi.org/10.1016/j.febslet.2014.03.055

25. Bashir Q, Volkov AN, Ullmann GM, Ubbink M (2010) Visualization of the encounter ensemble of the transient electron transfer complex of cytochrome c and cytochrome c peroxidase. J Am Chem Soc. https://doi.org/10.1021/ja9064574

26. Kim DE, Chivian D, Baker D (2004) Protein structure prediction and analysis using the Robetta server. Nucleic Acids Res. https://doi.org/10.1093/nar/gkh468

27. Webb B, Sali A (2016) Comparative protein structure modeling using MODELLER. Curr Protoc Bioinforma. https://doi.org/10.1002/cpbi.3

28. Yagi H, Pilla KB, Maleckis A, et al (2013) Three-dimensional protein fold determination from backbone amide pseudocontact shifts generated by lanthanide tags at multiple sites. Structure. https://doi.org/10.1016/j.str.2013.04.001

29. Volkov AN, Worrall JAR, Holtzmann E, Ubbink M (2006) Solution structure and dynamics of the complex between cytochrome c and cytochrome c peroxidase determined by paramagnetic NMR. Proc Natl Acad Sci. https://doi.org/10.1073/pnas.0603551103

30. Battiste JL, Wagner G (2000) Utilization of site-directed spin labeling and high-resolution heteronuclear nuclear magnetic resonance for global fold determination of large proteins with limited nuclear overhauser effect data. Biochemistry. https://doi.org/10.1021/bi000060h

31. Leaver-Fay A, O’Meara MJ, Tyka M, et al (2013) Scientific benchmarks for guiding macromolecular energy function improvement. In: Methods in Enzymology

32. Abraham MJ, Murtola T, Schulz R, et al (2015) Gromacs: High performance molecular simulations through multi-level parallelism from laptops to supercomputers. SoftwareX. https://doi.org/10.1016/j.softx.2015.06.001

33. Touw WG, Baakman C, Black J, et al (2015) A series of PDB-related databanks for everyday needs. Nucleic Acids Res. https://doi.org/10.1093/nar/gku1028

34. Rodrigues JPGLM, Trellet M, Schmitz C, et al (2012) Clustering biomolecular complexes by residue contacts similarity. Proteins Struct Funct Bioinforma. https://doi.org/10.1002/prot.24078

35. Keizers PHJ, Ubbink M (2011) Paramagnetic Tools in Protein NMR. In: Protein NMR Spectroscopy: Practical Techniques and Applications

36. Xu Q, Zhu S, Wang W, et al (2009) Regulation of kinetochore recruitment of two essential mitotic spindle checkpoint proteins by Mpsl phosphorylation. Mol Biol Cell. https://doi.org/10.1091/mbc.E08-03-0324

37. Hiruma Y, Sacristan C, Pachis ST, et al (2015) Competition between MPS1 and microtubules at kinetochores regulates spindle checkpoint signaling. Science (80-). https://doi.org/10.1126/science.aaa4055

38. Liu X, Winey M (2012) The MPS1 family of protein kinases. Annu Rev Biochem 81:561–585. https://doi.org/10.1146/annurev-biochem-061611-090435

39. Combes G, Barysz H, Garand C, et al (2018) Mps1 Phosphorylates Its N-Terminal Extension to Relieve Autoinhibition and Activate the Spindle Assembly Checkpoint. Curr Biol. https://doi.org/10.1016/j.cub.2018.02.002

